# Evaluation of HD-sEMG descriptor sensitivity to changes of anatomical and neural properties with aging : A simulation study

**DOI:** 10.1101/2024.09.26.612400

**Authors:** Ines Douania, Jeremy Laforet, Sofiane Boudaoud

## Abstract

**Background and objectives:** A reliable evaluation of anatomical and neural muscle properties and its effects on the electrical signals measured at the skin surface aims to develop a medical non-invasive aid-diagnosis tool assisted by model and personalized to the patient. This tool will be dedicated to understand and evaluate muscle diseases and aging.

**Methods:** We perform a new Robust Morris Screening Method: RMSM in Douania et al. (2023), to assess the impact of muscle anatomy (model inputs) uncertainties and variations on a simulated HD-sEMG signals (model outputs). The model describes a complex neuromuscular system simulating HD-sEMG (high density surface electromyography) signals generated from motor units electrical sources of a striated muscle: the Biceps Brachii (BB). Two subjects categories and two contractions levels are studied: young men (YM) and old men (OM) at low and high contraction (LC = 20% of MVC and HC = 60% of MVC). A 33 features in time and frequency domains are used as model outputs.

**Results:** We have demonstrated that the neuromuscular model is able to deliver HD-sEMG signals sensitive to the same anatomical and neural muscle factors as in real cases. Time domain features are mainly sensitive to muscle thickness, conduction velocity of fibers, electrode locations (at HC), number of motor units, and to the number of slow and fast fibers for young and aged categories respectively. Frequency domain feature are sensitive mainly to the conduction velocity of fibers and muscle conductivities (no significant differences between YM and OM are observed).

**Conclusion:** This result is important, it allows to obtain simulated HD-sEMG signals close to experimental ones with low cost and in reduced time. However, for a reliable evaluation of muscle aging, the neuromuscular model should be enhanced to better describe structural, morphological, and functional age-related phenomena.

## Introduction

Muscle aging is a progressive physiological process. It starts from the adult age (*∼*40 years), and involves a progressive loss of muscle mass and strength. Age-related muscle changes are driven by a combination of many phenomena and factors affecting the composition, the structure, the morphology and the functions of muscle tissues. The muscle composition alterations can be a metabolic disorder and/or a protein turnovers due to a multiple factors, e.g., insulin resistance, hypertension, or genetic factors Nair (2005). The muscle structural and morphological changes are displayed by a decrease of fiber sizes Larsson (1978); Verdijk et al. (2007) and number Lexell et al. (1983); Sjöström et al. (1992), and a different apportionment of slow and fast fibers within muscle. The decline of muscle functions is a consequence pronounced by previous mentioned phenomena and leading to a progressive muscle force reduction. A high decline rate of these aging changes can induce critical diseases: e.g., sarcopenia and cancer cachexia Tieland et al. (2018).

Factors influencing this rate can be detected by surface electromyography signals (sEMG) which is an essential non-invasive tool to evaluate muscle activation and strength in biomedical diagnostics. It is used to detect, monitor, and predict age-related alterations of the neuromuscular system Boccia et al. (2015).

The current study is motivated by identifying most influential parameter(s) and muscle aging phenomena of neuro-muscular system on the recorded sEMG signal, for young and elder people through a recent Robust Morris Screening Method. This identification will allow to angle the experimental effort toward the most important age-related changes. In addition, it will differentiate and retrieve important neuromuscular parameters, not easily achievable, to identify by inverse method using sEMG signal. For this purpose, we will use a HD-sEMG fast generation model Carriou et al. (2016, 2018), simulating motor units action potential (MUAP) at the skin surface. In this model, the MUAPs depend of muscle and conductor volume anatomy. More than 50 parameters (Fig. 4) are involved in this model. Parameters describe the current sources (sizes, locations, types), muscle and conductor volume morphology and structure, and the 64 HD-sEMG recording system. Using variation ranges of these parameters reported from literature, we define two age categories: young men (YM) and elder men (EM). The assessment of influential parameters for each category will be deployed at two contraction levels:(i) Low contractions (LC) at 20% of maximum voluntary contraction (MVC), and (ii) High contractions (HC) at 60% of MVC. A 33 amplitude, frequency and statistical features of HD-sEMG simulated signals are used to identify these parameters and quantifying/qualify its impacts.

**Figure 1.**
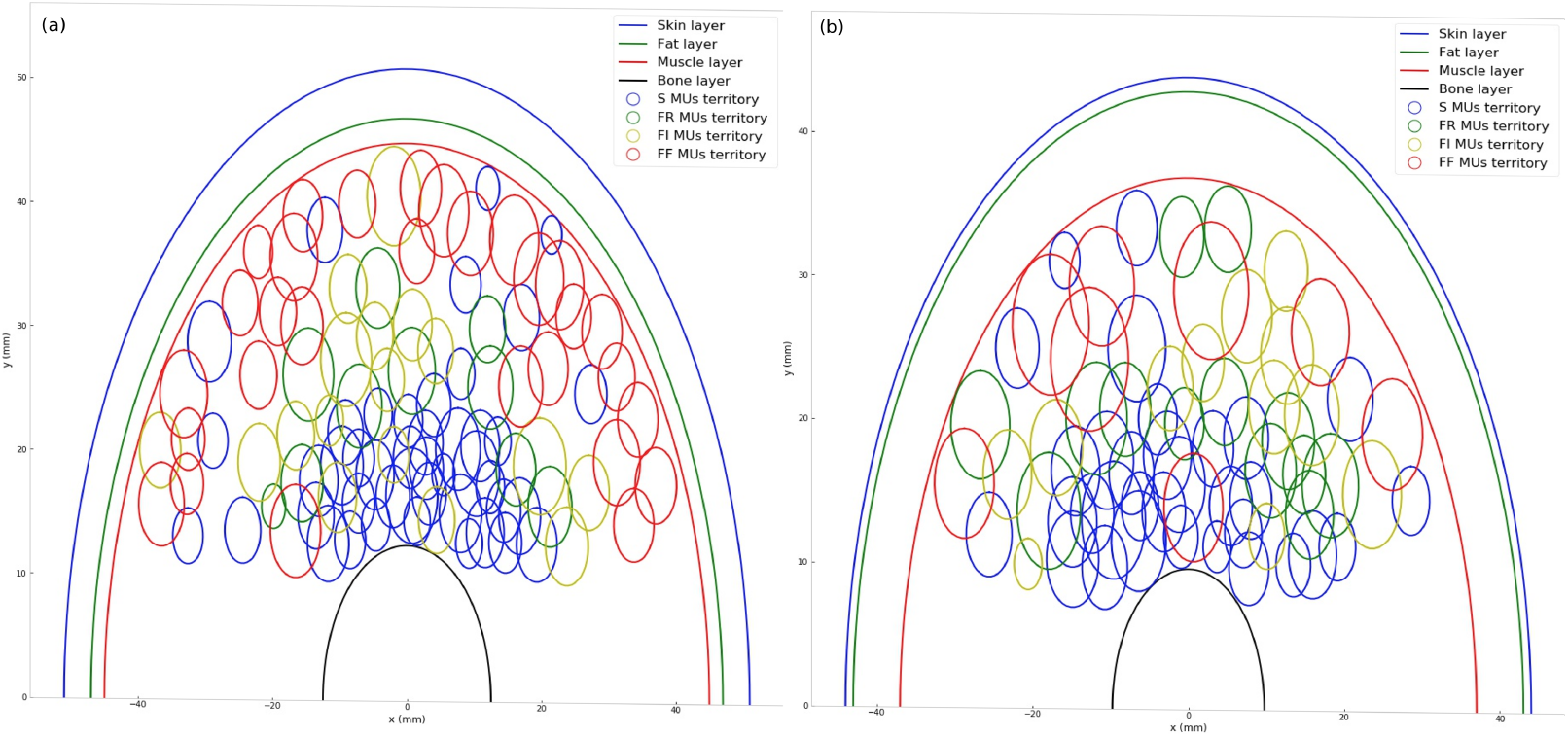
Source locations within a cylindrical muscle volume. (a) Young muscle. (b) Aged muscle

**Figure 2.**
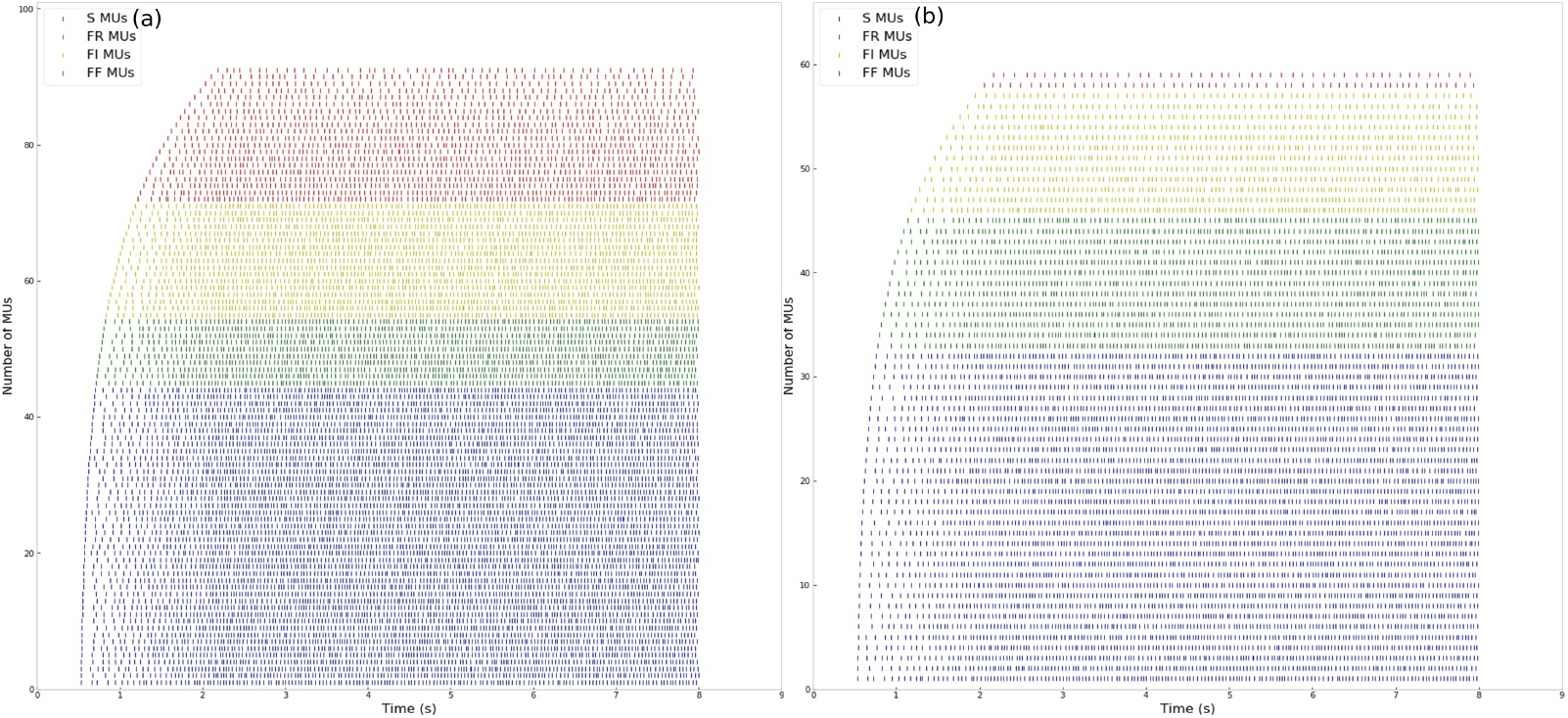
Recruitment Pattern. (a) Young muscle. (b) Aged muscle

**Figure 3.**
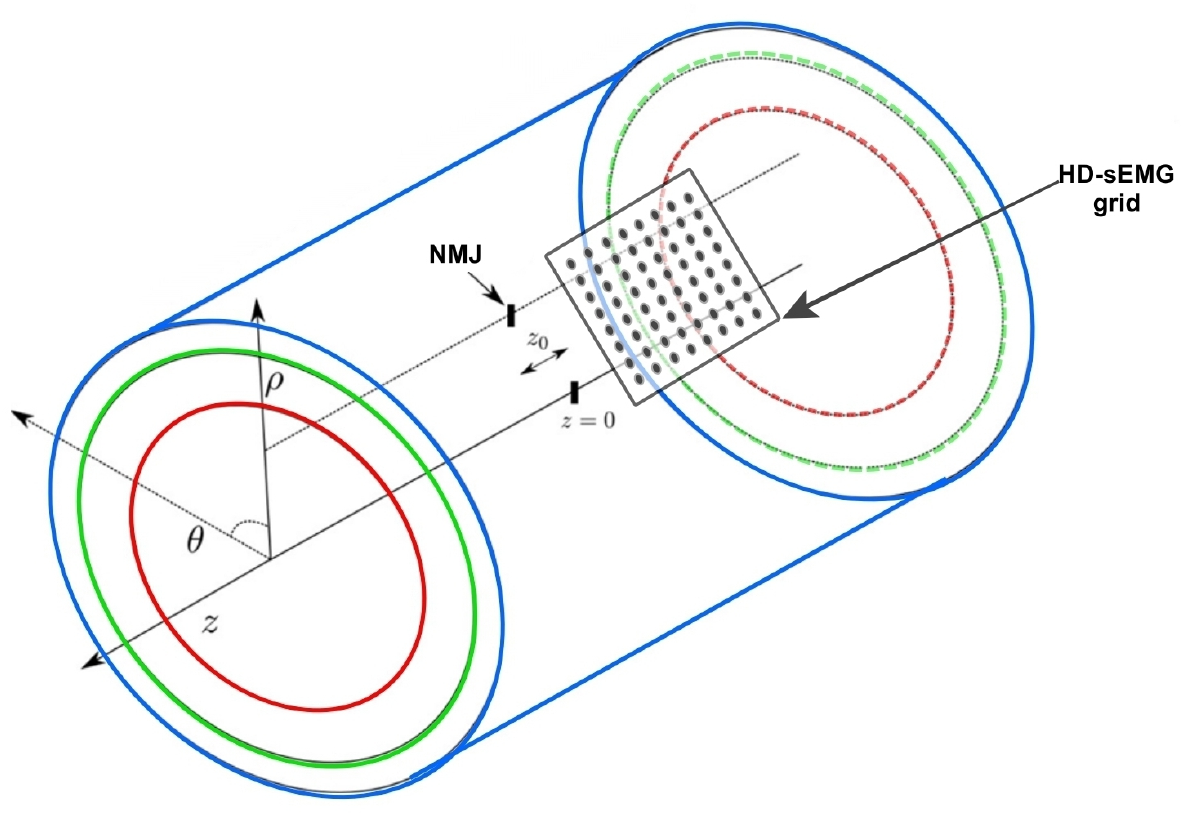
Muscle model geometry and position of the recording system.

**Figure 4.**
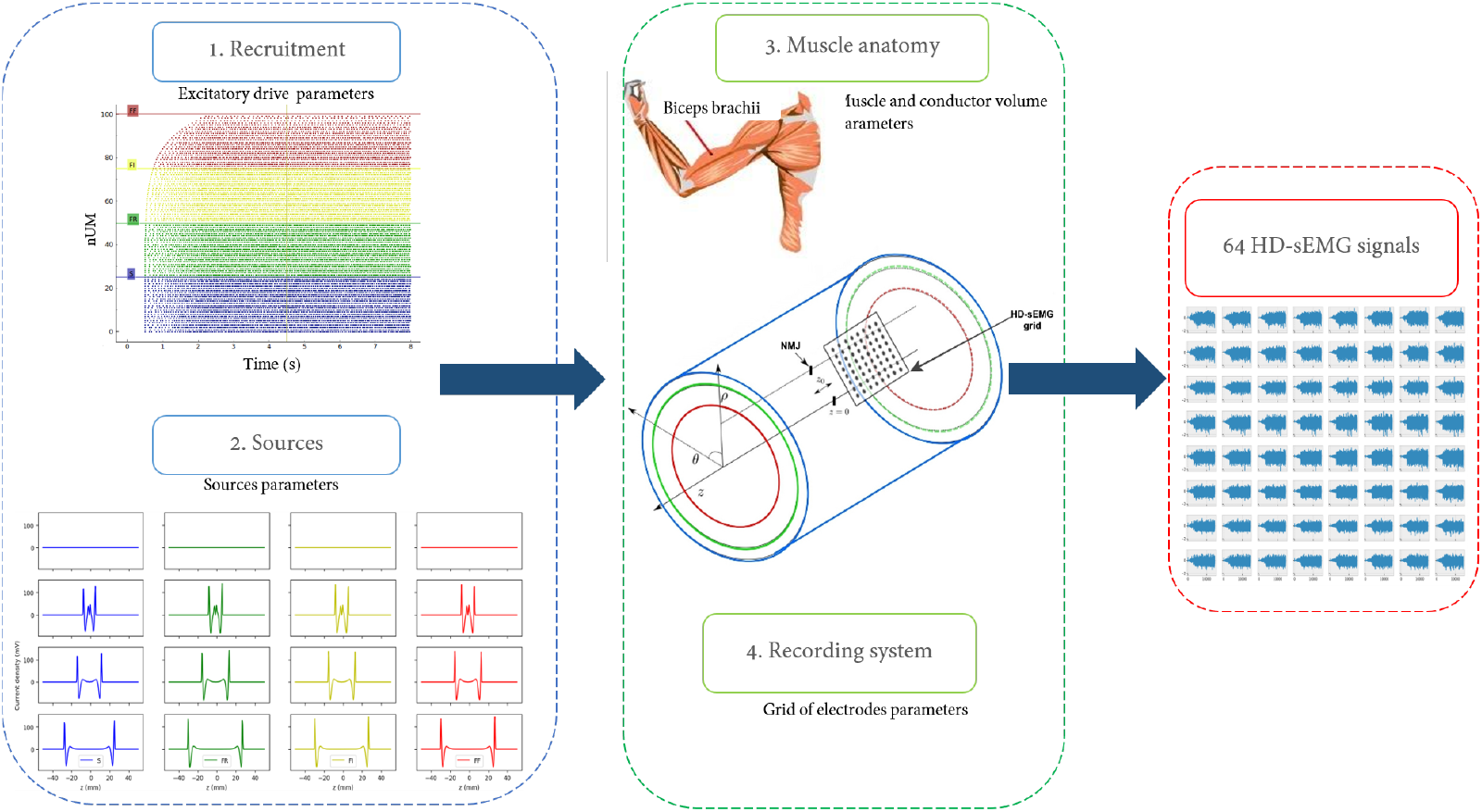
Modeling approach.

It was demonstrated that sEMG amplitude during sub-MVC is lower for elderly than young adults. With the frequency spectrum analysis, the signal frequency mean is higher when faster motor units (MUs) are recruited. During aging, this mean/median shows a marked decreasing rate for elderly subjects at isometric contractions (predominant slow fiber muscle González-Izal et al. (2012); Macaluso and De Vito (2004)) More influential co-factors can be related to this decrease (e.g., fiber length, depth of MU within the muscle, or volume conductor layers. Farina (2008)). Moreover, many advanced techniques are used to evaluate muscle aging by estimating from sEMG the: (i) fiber conduction velocity which increases when faster and larger fibers are recruited (young population muscles), and (ii) MUs attributes (number, size, recruitment thresholds, firing rate).

The most previous cited techniques and methods to extract information from sEMG allow a good differentiation between populations studied according to the age. However, these methods cannot differentiate and quantify the impact of each age-related characteristic/phenomena on the sEMG (e,g,. atrophy, fiber apoptosis). Moreover, using advanced techniques, we can estimate some neuromuscular factors related to these phenomena (e,g,. MUs number, fiber conduction velocity and size) but are not able to relate changes observed on sEMG signal between young and elder population to a precise anatomical or/and neural muscle factor or group of factors. Studies cannot compare impacts related to these age-related changes using only sEMG.

In this study, we present an improved sensitivity analysis applied on a realistic bio-reliable simulator of HD-sEMG signals from a Biceps brachii (BB) muscle during isometric contractions. The task is done by exploring 33 features of HD-sEMG, achieved from a realistic young and elder muscle anatomies.

## MATERIALS AND METHODS

In this study, a robust and reliable screening method (RMSM) is applied Douania et al. (2023). The RMSM method provides a global sensitivity analysis with a reduced number of model evaluations. In this paper, we will (i) first briefly summarize the modeling approach as depicted in Carriou et al. (2016): inputs and features (outputs), then (ii) we will describe the RMSM method: strategy and indices, and finally (iii) we will present the simulation plan to perform the RMSM results in section D: age categories and contraction levels.

### A. Modeling approach

The modeling approach depicted in Carriou et al. (2016, 2018) simulates the electrical activity of BB muscle during isometric contractions (Fig. 4). The main axis of this neuromuscular model are:

1. The modeling of the sources (MUs): Fours types of MUs are displayed with circular territories (Slow (S), Fast Intermediate (FI), Fast Resistant (FR) and Fast Fatigable (FF)). Faster MUs are placed closer to muscle surface and slower ones are located in deeper muscle layers. Fibers are defined (types and positions) based on the corresponding MUs.
2. The recruitment and firing pattern: MUs are activated according to specific threshold and its recruitment is regulated by force dependent firing rate.
3. The conductor volume: a multilayered cylindrical volume is used (skin, adipose tissues, and muscle tissues). Sources are modeled at the microscopic muscle scale and its intracellular potential is generated and propagation along its fibers.
4. The recording system: the sEMG grid is placed at the most external boundary of the conductor volume (skin layer). The sEMG signal for each electrode is modeled as an electrode surface integration of electrical activity values over the sampled positions under this surface. the configuration of electrodes can be adjustable (location, number, and shape).

In this model, computation of the sEMG is fully performed in a three dimensional Fourier domain. Details about the resolution of the Poisson equation and computation in Fourier domain are provided in Carriou et al. (2016).

#### A.1. Model inputs

In this study, we investigate sensitivity of HD-sEMG signals to the variation of 35 parameters of the neuromuscular model. Variation intervals are extracted from literature respecting a specific considerations:

- Intervals are extracted from BB muscle, during isometric contractions, and for healthy young and old men.
- Intervals are selected based on the methods and techniques involved in the measurement, and the clarity of information related to the age and gender of subjects in material and methods paragraphs.
- Priority was assigned to values extracted by autopsy or biopsy for some parameters like the fibers number and length. Priority is also given to recent methods of measurement (e.g., Magnetic Resonance Imaging) to define parameters like bone radius or muscle thickness.
- For parameters not easily achieved using previous techniques like the MU distribution per type or its positions, the selection is based on the reliability of three essential factors: the performance of the recording system; the experimental protocol used and if it is adequate with our criteria (e.g., isometric contraction); the reliability of the algorithms applied to extract data from recorded signal.

#### A.2. Model outputs

The main output of the model is the HD-sEMG signal reflecting, through complex analysis methods, many physiological and anatomical properties of the neuromuscular system. In order to correlate this signal to these properties, many features are reported in the literature. Feature extraction is the first and crucial step in the signal processing. Table 2 summarize all the features extracted from the HD-sEMG signal and considered as the model outputs for this study. Features depicted in Table 2 can be classified according to their computation algorithm techniques (Fig. 5). We have classified features into mono-variate and bi-variate subgroups, according to the fact of using one electrode channel or two. Then, the mono-variate features are classified either as time, frequency domain or non-linear based algorithms.

**Table 1.** List of parameters with variation ranges extracted from literature. Values for young men (YM) and old men (OM). *N* normal distribution. *U* uniform distribution. S: Slow, FI: Fast Intermediate, FR: Fast Resistant, FF: Fast Fatiguable. (−) No measurement unit.

| Name | Description | Unit | YM value | OM value |
| --- | --- | --- | --- | --- |
| $n_{MU}$ | Number Motoneuron Unit | - | $\mathcal{U}[350, 450]$ | $\mathcal{U}[250, 350]$ |
| $MU_{distribution}$ | Distribution of MU according to their type | (%) | $\mathcal{N}_S(47, 8)$<br>$\mathcal{N}_{FI}(10, 11)$<br>$\mathcal{N}_{FR}(20, 6)$<br>$\mathcal{N}_{FF}(29, 11)$ | $\mathcal{N}_S(52, 8)$<br>$\mathcal{N}_{FI}(18, 2)$<br>$\mathcal{N}_{FR}(15, 6)$<br>$\mathcal{N}_{FF}(12, 9)$ |
| $MU_S^r$ | Radius of MU type S | mm | $\mathcal{N}(2.5, 0.5)$ | $\mathcal{N}(2.75, 0.5)$ |
| $MU_{FI}^r$ | Radius of MU type FI | mm | $\mathcal{N}(2.75, 0.5)$ | $\mathcal{N}(3.0, 0.5)$ |
| $MU_{FR}^r$ | Radius of MU type FR | mm | $\mathcal{N}(3, 0.5)$ | $\mathcal{N}(3.25, 0.5)$ |
| $MU_{FF}^r$ | Radius of MU type FF | mm | $\mathcal{N}(3.25, 0.5)$ | $\mathcal{N}(3.5, 0.5)$ |
| $n_{SMU}^f$ | Number of fibers per MU type S | mm | $\mathcal{N}(240, 310)$ | $\mathcal{U}(160, 210)$ |
| $n_{FIMU}^f$ | Number of fibers per MU type FI | - | $\mathcal{N}(80, 100)$ | $\mathcal{U}(80, 90)$ |
| $n_{FRMU}^f$ | Number of fibers per MU type FR | - | $\mathcal{N}(100, 150)$ | $\mathcal{U}(100, 140)$ |
| $n_{FFMU}^f$ | Number of fibers per MU type FF | - | $\mathcal{N}(170, 260)$ | $\mathcal{U}(120, 200)$ |
| $L$ | Muscle length | mm | $\mathcal{U}(115, 130)$ | $\mathcal{U}(95, 115)$ |
| $bone_r$ | Bone radius | mm | $\mathcal{U}(11.7, 13.2)$ | $\mathcal{U}(9.5, 13)$ |
| $S_{fD}$ | S fiber diameter | $\mu m$ | $\mathcal{N}(66, 9)$ | $\mathcal{N}(51, 6)$ |
| $FI_{fD}$ | FI fiber diameter | $\mu m$ | $\mathcal{N}(73, 15)$ | $\mathcal{N}(56, 13)$ |
| $FR_{fD}$ | FI fiber diameter | $\mu m$ | $\mathcal{N}(76, 15)$ | $\mathcal{N}(60, 11)$ |
| $FF_{fD}$ | FF fiber diameter | $\mu m$ | $\mathcal{N}(74, 15)$ | $\mathcal{N}(51, 15)$ |
| $C_{velocity}$ | Conduction velocity | $m.s^{-1}$ | $\mathcal{U}[3.5, 5.9]$ | $\mathcal{U}[2.5, 4.2]$ |
| $NMJ_{pos}$ | Neuromuscular junction center | - | $\mathcal{U}[-15, 15]$ | $\mathcal{U}[-15, 15]$ |
| $MTZ_L$ | Left myotendinous length | mm | $\mathcal{N}(15, 2)$ | $\mathcal{N}(15, 2)$ |
| $MTZ_R$ | Right myotendinous length | mm | $\mathcal{N}(15, 2)$ | $\mathcal{N}(15, 2)$ |
| $\rho_{muscle}^c$ | Radial muscle conductivity | $S.m^{-1}$ | $\mathcal{U}[0.4, 0.6]$ | $\mathcal{U}[0.3, 0.56]$ |
| $\theta_{muscle}^c$ | Angular muscle conductivity | $S.m^{-1}$ | $\mathcal{U}[0.4, 0.6]$ | $\mathcal{U}[0.3, 0.5]$ |
| $Z_{muscle}^c$ | Longitudinal muscle conductivity | $S.m^{-1}$ | $\mathcal{U}[0.81, 0.99]$ | $\mathcal{U}[0.73, 0.96]$ |
| $thick_{muscle}$ | Muscle thickness | mm | $\mathcal{N}(44, 2)$ | $\mathcal{N}(40, 2)$ |
| $\rho_{Fat}^c$ | Radial fat conductivity | $S.m^{-1}$ | $\mathcal{U}[0.04, 0.07]$ | $\mathcal{U}[0.04, 0.07]$ |
| $\theta_{Fat}^c$ | Angular fat conductivity | $S.m^{-1}$ | $\mathcal{U}[0.04, 0.07]$ | $\mathcal{U}[0.04, 0.07]$ |
| $Z_{Fat}^c$ | Longitudinal fat conductivity | $S.m^{-1}$ | $\mathcal{U}[0.04, 0.07]$ | $\mathcal{U}[0.04, 0.07]$ |
| $thick_{Fat}$ | Fat thickness | mm | $\mathcal{U}[4, 6]$ | $\mathcal{U}[4, 7]$ |
| $\rho_{skin}^c$ | Radial skin conductivity | $S.m^{-1}$ | $\mathcal{U}[0.9, 1.2]$ | $\mathcal{U}[0.9, 1.2]$ |
| $\theta_{skin}^c$ | Angular skin conductivity | $S.m^{-1}$ | $\mathcal{U}[0.9, 1.2]$ | $\mathcal{U}[0.9, 1.2]$ |
| $Z_{skin}^c$ | Longitudinal skin conductivity | $S.m^{-1}$ | $\mathcal{U}[0.9, 1.2]$ | $\mathcal{U}[0.9, 1.2]$ |
| $thick_{skin}$ | Skin thickness | mm | $\mathcal{U}[0.9, 1.2]$ | $\mathcal{U}[0.9, 1.2]$ |
| $Grid_\theta$ | Angular grid center | $^\circ$ | $\mathcal{U}[0, 5]$ | $\mathcal{U}[0, 5]$ |
| $Grid_z$ | Longitudinal grid center | mm | $\mathcal{U}[22, 26]$ | $\mathcal{U}[22, 26]$ |
| $Grid_{rot}$ | Electrode grid rotation | $^\circ$ | $\mathcal{U}[0, 10]$ | $\mathcal{U}[0, 10]$ |

**Table 2.** List of Features extracted from the HD-sEMG and considered as the model outputs.

| Feature | Abbreviation |
| --- | --- |
| 1.Mean Absolute Value | MAV |
| 2.Modified Mean Absolute Value 1 | MAV1 |
| 3.Modified Mean Absolute Value 2 | MAV2 |
| 4.Integrated EMG | IEMG |
| 5.Sample Square Integrated | SSI |
| 6.Waveform Length | WL |
| 7.Average Amplitude Change | AAC |
| 8.Difference Absolute Standard Deviation Value | DASDV |
| 9.Root Mean Square of Amplitude | RMSA |
| 10. Average Rectified Value | ARV |
| 11.Variance of EMG | Var |
| 12.Mean of EMG | Mean |
| 13.Myopluse percentage rate | Myopulse rate |
| 14.Number of Zero Crossing of EMG | NoZC |
| 15.Wilson Amplitude | wilson_amp |
| 16.Kurtosis | Kurt |
| 17.Skewness | Skew |
| 18.Mean power | MNP |
| 19.Total power | TTP |
| 20.Median frequency | MDF |
| 21.Mean frequency | MNF |
| 22.Peak frequency | Peak_f |
| 23.Power spectrum ratio | Power_r |
| 24.Frequency ratio | Freq_r |
| 25.Spectral moment 1 | SM1 |
| 26.Spectral moment 2 | SM2 |
| 27.Spectral moment | SM3 |
| 28.Variance of central frequency of PSD | VDF |
| 29.Time reversibility | Trev |
| 30.Approximate entropy | ap_ent |
| 31.Sample entropy | sp_ent |
| 32.Non-linear correlation coefficient | $H_2P$ |
| 33.Pearson correlation coefficient | $R_2P$ |

**Figure 5.**
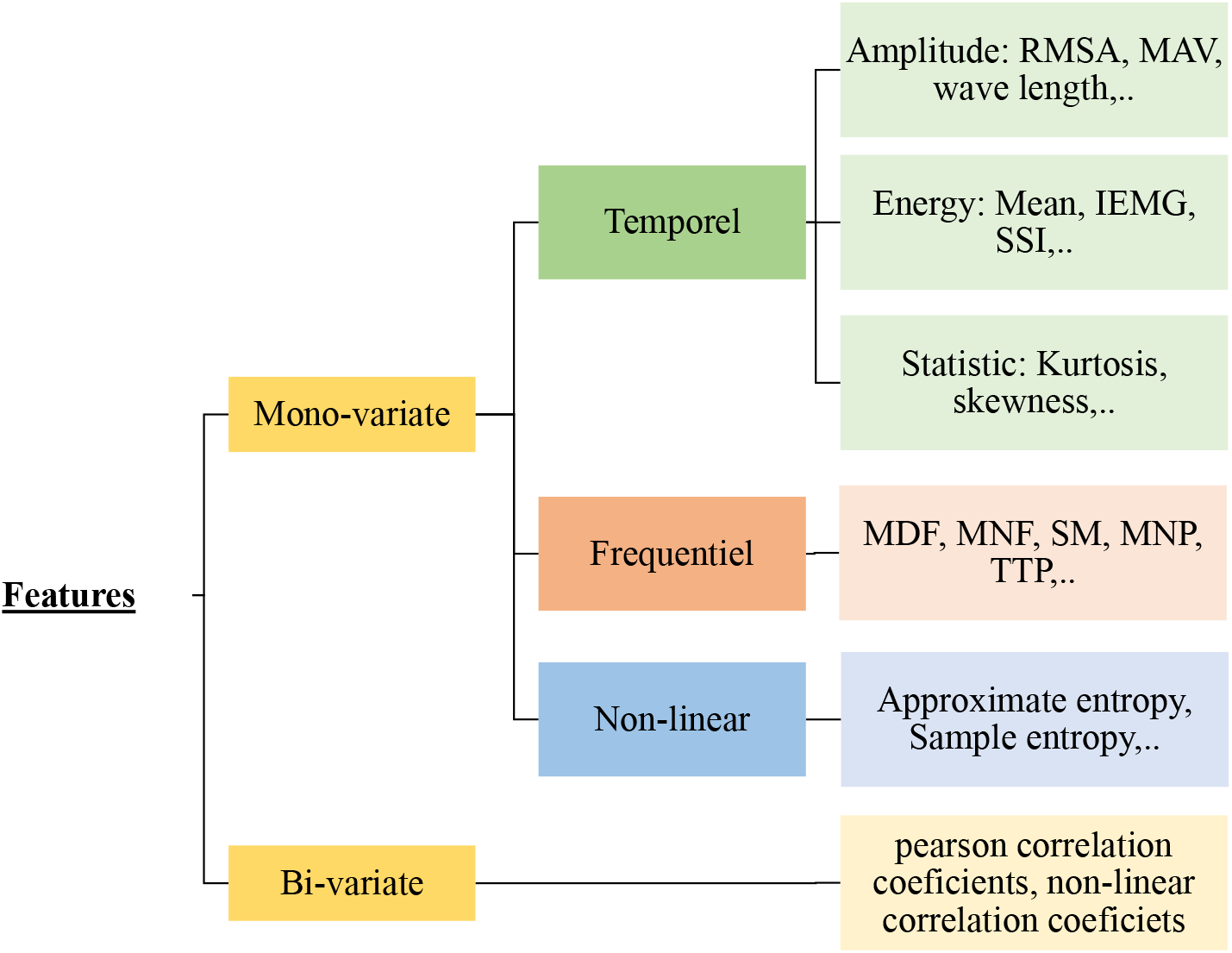
Classification of HD-sEMG features.

### B. Sensitivity analysis

The neuromuscular model inputs have : (i) a large number (35 parameters are selected from 50), (ii) a large variation ranges, and (iii) a high degree of interaction. Furthermore, the model is a large and complex system, with high computation time (CT). One simulation with *nUM* = 400 and at 60% of maximal volunteer contraction (MVC) turns in around 40 minutes. As consequence, methods can performing a reliable sensitivity analyses with reasonable CT are limited. Variance based global sensitivity analysis methods are eliminated. We perform a screening sensitivity analysis method (RMSM). The Table 3 shows the estimated CT for the RMSM and Sobol (a variance-based method Sobol (2001)) methods with only 35 model inputs. Parameters related to recruitment pattern and some electrode grid characteristics (e.g, diameter of electrodes) are fixed. The Table 3 shows an acceptable CT (*∼* 89h) RMSM. In contrast, we will need 374 days to obtain a sensitivity analysis with the Sobol method. A deeper SA analysis with variance-based method can be performed on a reduced sample of model inputs identified after applying RMSM method.

**Table 3.** Computation time with RMSM and Sobol methods. Model with: N = number of parameters = 35; *nUM* = [120, 400]; and contractions level = 70%MVC. Number of trajectories for RMSM = T = 10.

| Machine configuration | Method | Number of evaluation | Total computation time |
| --- | --- | --- | --- |
| 2 x 8 core intel Xeon 2.4 GHz with hyper threading (32 threads)<br>128 Go RAM | RMSM | $\approx T \times N + 1 \approx 380$ | 14h28mn (89h09mn 1 CPU machine) |
| | Sobol | $\sim 1000 \times N \approx 37000$ | 59Days5h (374Days5h with 1 CPU machine) |

### C. RMSM method

The RMSM evaluates the distribution of the elementary effects (EE) of each input factor on the *k*^*th*^ model output as depicted in Morris (1991), from which a basic statistics (average and deviation) are computed to derive sensitivity information. One elementary effect of the input *X*_*j*_ on the output *y*_*k*_ is computed as following:

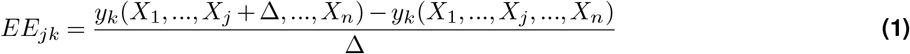

Where, Δ is a predetermined perturbation factor of *X*_*j*_, *y*_*k*_(*X*_1_, *X*_2_, …, *X*_*j*_, …, *X*_*n*_) is the scalar model output evaluated at input parameters (*X*_1_, *X*_2_, …, *X*_*j*_, …, *X*_*n*_), while *y*_*k*_(*X*_1_, *X*_2_, …, *X*_*j*_ + Δ, …, *X*_*n*_)is the scalar output corresponding to a Δ changes in *X*_*j*_. Each input parameter *X*_*j*_ can only take values corresponding to predefined set of p levels from its variation range. The computation of EE is repeated *T* times also named trajectory. Thus, the designation of “one factor at a time” (OAT) of this screening method. This design requires *T ∗* (*n* + 1) model evaluations. Each EE distribution due to the *j*_*th*_ input variable on the *k*_*th*_ output contains *T* independent elementary effects. Indices assessing the impact of inputs are the absolute median and the median absolute deviation computed from EE distribution. The following equations describe the RMSM indices:

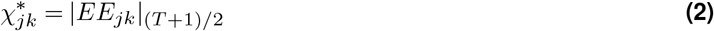

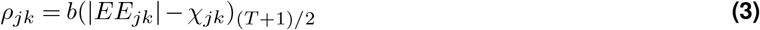

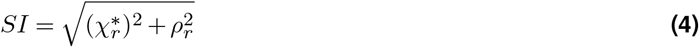

### D. Simulations plan and results exposure design

#### simulations plan

Two categories of subjects are studied: Young Men (YM) and Old Men (OM). Two contraction levels are considered: Low Contractions (20% of MVC) and High Contractions (60% of MVC). The total number of sensitivity analysis performed is eight (Fig. 6).

**Figure 6.**
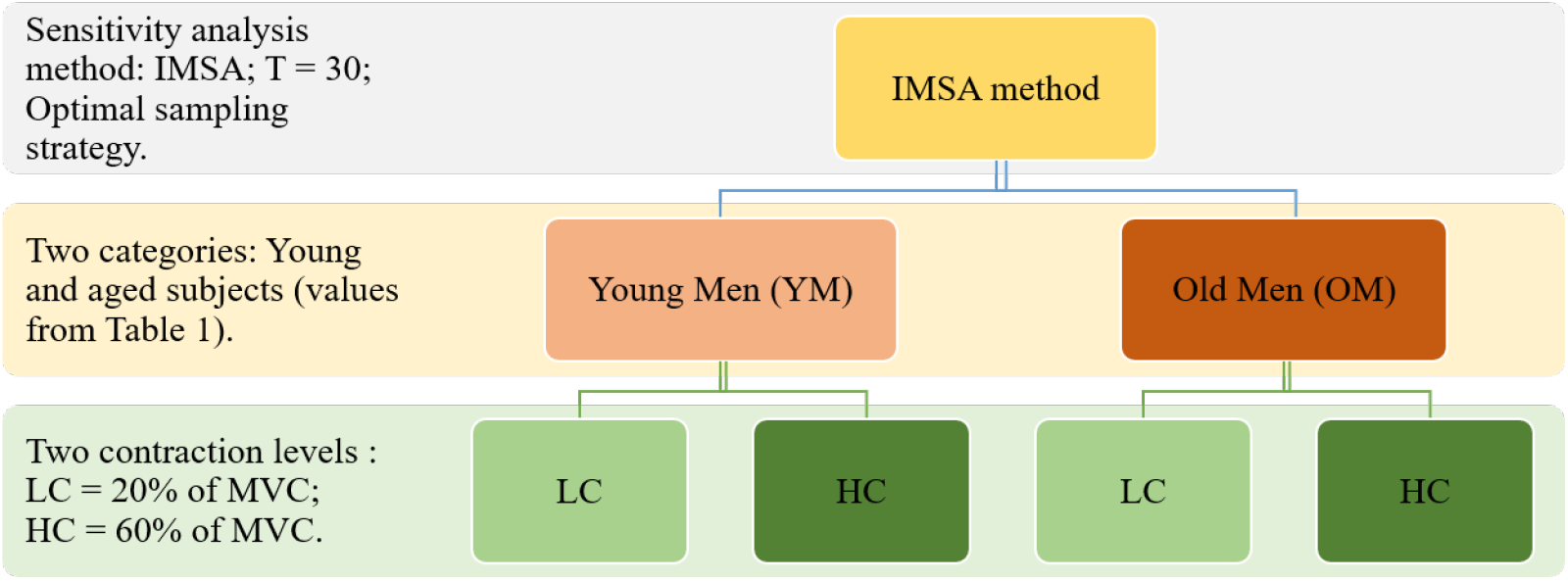
Simulation plan. Four sensitivity analysis: YM at LC; YM at HC; OM at LC; OM at HC

#### Presentations of results

Results are presented mainly as cluster maps with min/max normalized *SI* indices (between 0 and 1). For each subgroup of outputs, each age category, and for each contractions level, a cluster map gives a relationship map between inputs and outputs.

#### Extraction of information towards parameters identification

We expect to have high colinearity among at least some of the computed features as they are close in concepts. To select the most informative subset, we decided to iteratively eliminate the features showing the highest colinearity. To do so, we compute the Variance inflation factor (VIF) on all features, remove the feature with the highest VIF, and repeat until the highest VIF is under the threshold value of 10. This process will be applied to all 4 conditions defined in the previous paragraph.

Then the remaining feature will be used to score the parameters using classical methods from the literature implemented in the Python library **ranky**: Majority judgement, Borda count, Copeland’s method, Random dictator. These scores will enable us to select the most influential parameters regardless of a specific feature ranking.

## RESULTS

### E. Mono-variate features

The RMSM was applied on a large and multi scales neuromuscular system. The number of trajectories is fixed at *T* = 30 and optimal sampling strategy is adopted Douania et al. (2023); Campolongo et al. (2007). Two subject categories are studied: YM and OM. The evaluation of input impacts is performed at two contraction levels: LC = 20% of MVC and HC = 60% of MVC. The impact of each input is assessed with features of Table 2. Each feature is computed from a 64 HD-sEMG simulated signals. Parallel computation with 32 CPUs intel Xeon calculator is used.

#### E.1. Time domain and non-linear features

Eleven features from Table 2 are selected: four amplitude features (RMSA, MAV, MAV1, MAV2), Three energy features (IEMG, SSI, Variance), two statistic features (Kurtosis, Skewness), and two non-linear features (ap_entropy, samp_entropy). The rest of features in these categories show the same results.

##### Amplitude and energy features

Features of this subgroup are easy to implement with the capacity to assess the main force generated by muscle. They can estimate efficiently the amount of muscle activity and fatigue. Correlating anatomical muscle properties to muscle force and fatigue is important for clinical and research fields. We observe, in Fig. 7, that all categories (YM, OM) and contraction levels (LC, HC) share approximately the influential and non-influential inputs. However, few changes are noted between Fig. 7(a), (b), (c), and (d). For YM (a & b), the number of fast fibers per MU (*MU* _*FF* _*nbF ibers*) has an important impact at HC. Which is not the case at LC (negligible effect), where slow fibers *MU* _*S*_*nbF ibers* have more effects. Likewise, electrode grid properties (position and rotation: *Grid*_*Z*_, *Grid*_*rot*_), total MU number (*nMU*), and radial conductivity of muscle 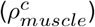 have more impact at HC. Its influences at LC is not negligible but not important. For OM, we note the same observation for slow and fast fibers between HC and LC. In addition, we observe that MU repartition is an important factor at HC and a negligible one at LC. The electrode grid properties have large impact at HC. Otherwise, OM share the same influential and non-influential parameters. Comparing changes between young and old categories, we cannot find differences at LC, expect a slight increase of bone radius (*bone*_*r*_) and muscle length (*L*) effects for OM. At HC, we observe same trends as LC, with the fact that (*MU*_*distribution*_)has large effect for OM.

**Figure 7.**
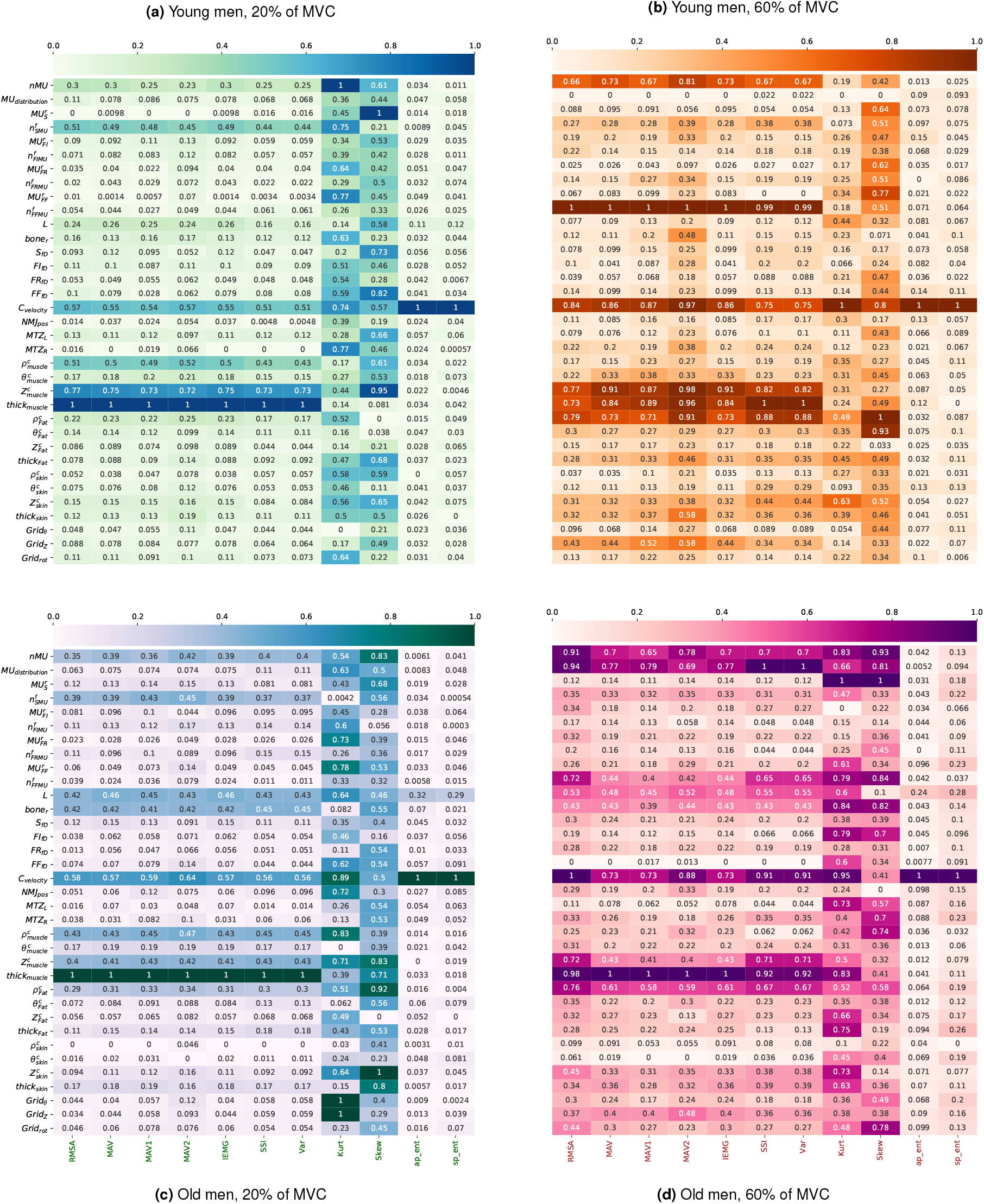
Impacts of model inputs on extracted amlitude HD-sEMG TD features (RMSM results). RMSM impact indice = Normalized *SI* (from 0 (light color) to 1 (dark color)). Ranking of neuromuscular inputs according to age (young (a,b) and old men (c,d)), and level of force contractions (low (a,c) and high(b,d)).

##### Statistic features

Kurtosis and skewness measure the peakedness and the symmetry/asymmetry of HD-sEMG signal respectively. The Fig. 7 shows that these two features have a different behaviors for each category. We cannot identify clearly a reduced and common number of influential inputs. The RMSM screening depicted in Fig. 8 (a, b) shows that all parameters have a close positions and situated at the non linear and/or with interactions effects zone, in contrast of amplitude and energy features (e.g., the RMSA feature in Fig. 8(c)). The stability of ranking is not assumed for these features due to the very close RMSM indices values. For that reason, we cannot find common trends between categories.

**Figure 8.**
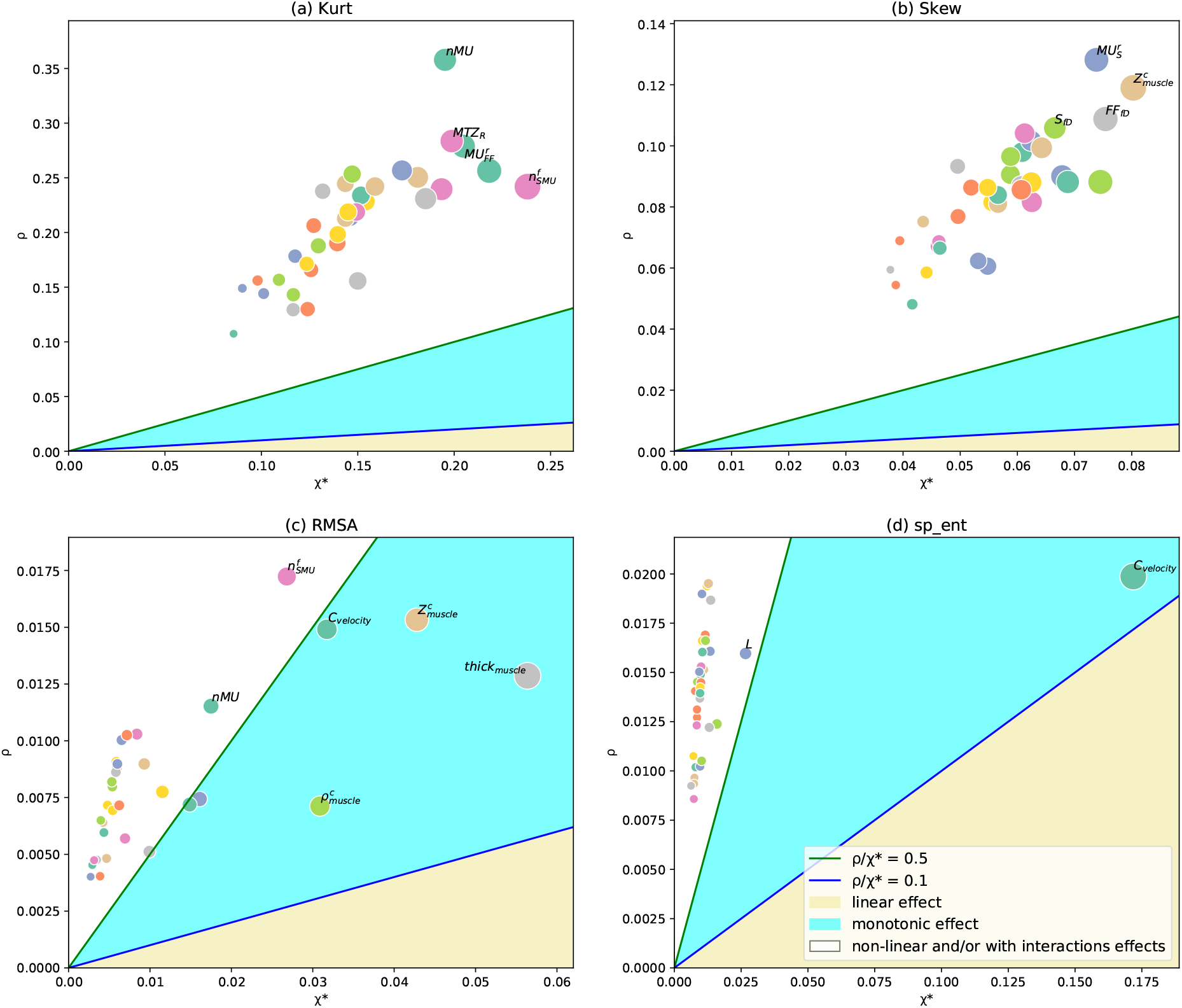
Parameters screening with RMSM method for (a) Kurtosis, (b) Skewness, (c) RMSA, and (d) sample entropy features. Categories: YM, LC.

##### Non linear features

The conduction velocity along fibers (*C*_*velocity*_) is the most influential parameter for entropy features: approximate entropy (*ap*_*ent*) and sample entropy (*sp*_*ent*) for both YM and OM at LC and HC. Entropy features are used to identify regularity and predictability of the signals. The conduction velocity has a large and monotonic effect on these features. The rest of of inputs have a negligible effects (Fig. 7 and Fig. 8(d)).

#### E.2. Frequency domain features

Frequency domain (FD) features are computed from Power Spectral Density (PSD). The welch method is performed to calculate PSD with “Hanning Windows”. These features required more CT compared to TD, and are useful in the detection of muscle fatigue. Ten FD features are computed (Fig. 9).

**Figure 9.**
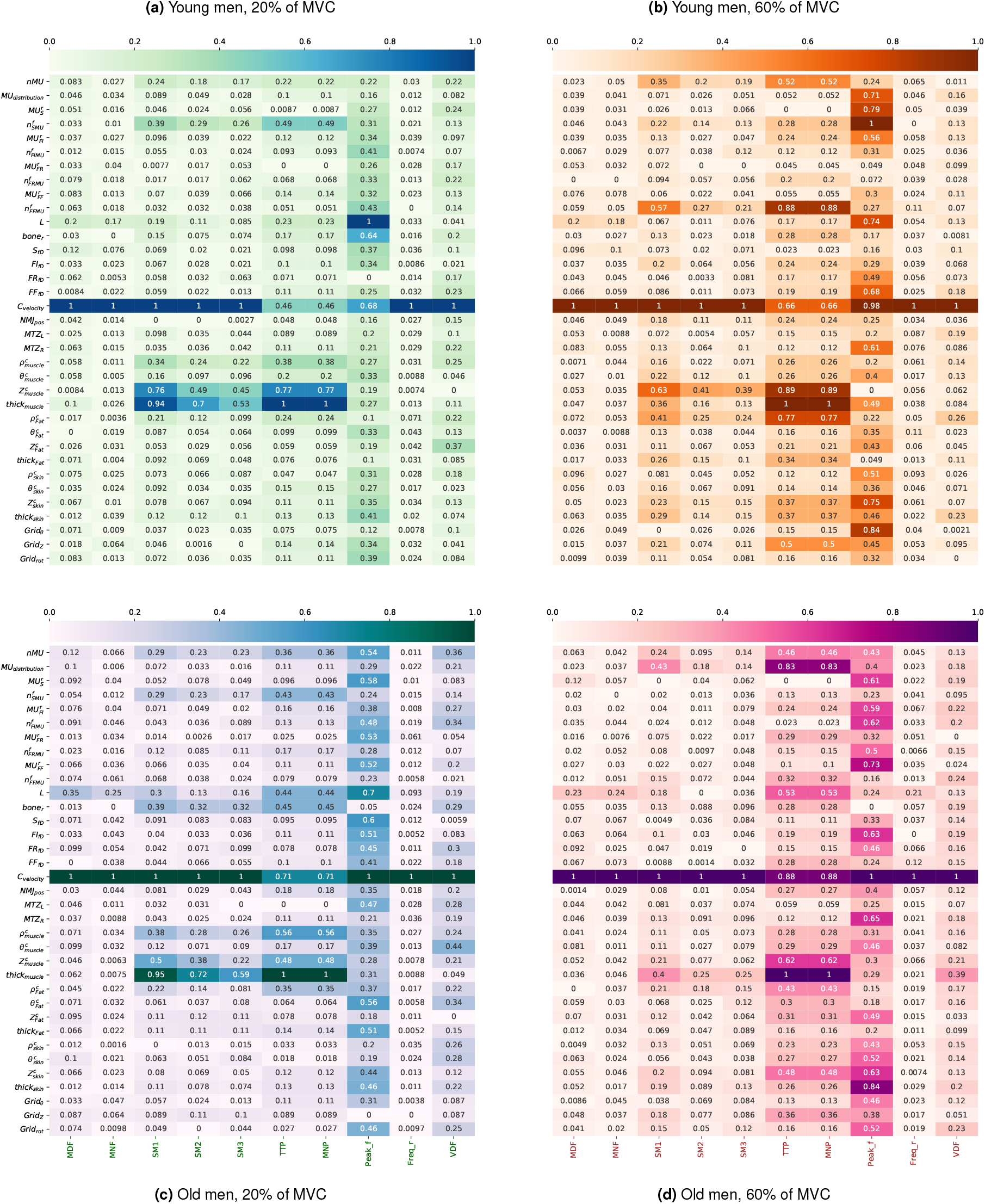
Impacts of model inputs on extracted amlitude HD-sEMG FD features (RMSM results). RMSM impact indice = Normalized *SI* (from 0 (light color) to 1 (dark color)). Ranking of neuromuscular inputs according to age (young (a,b) and old men (c,d)), and level of force contractions (ltheow (a,c) and high(b,d)).

**Figure 10.**
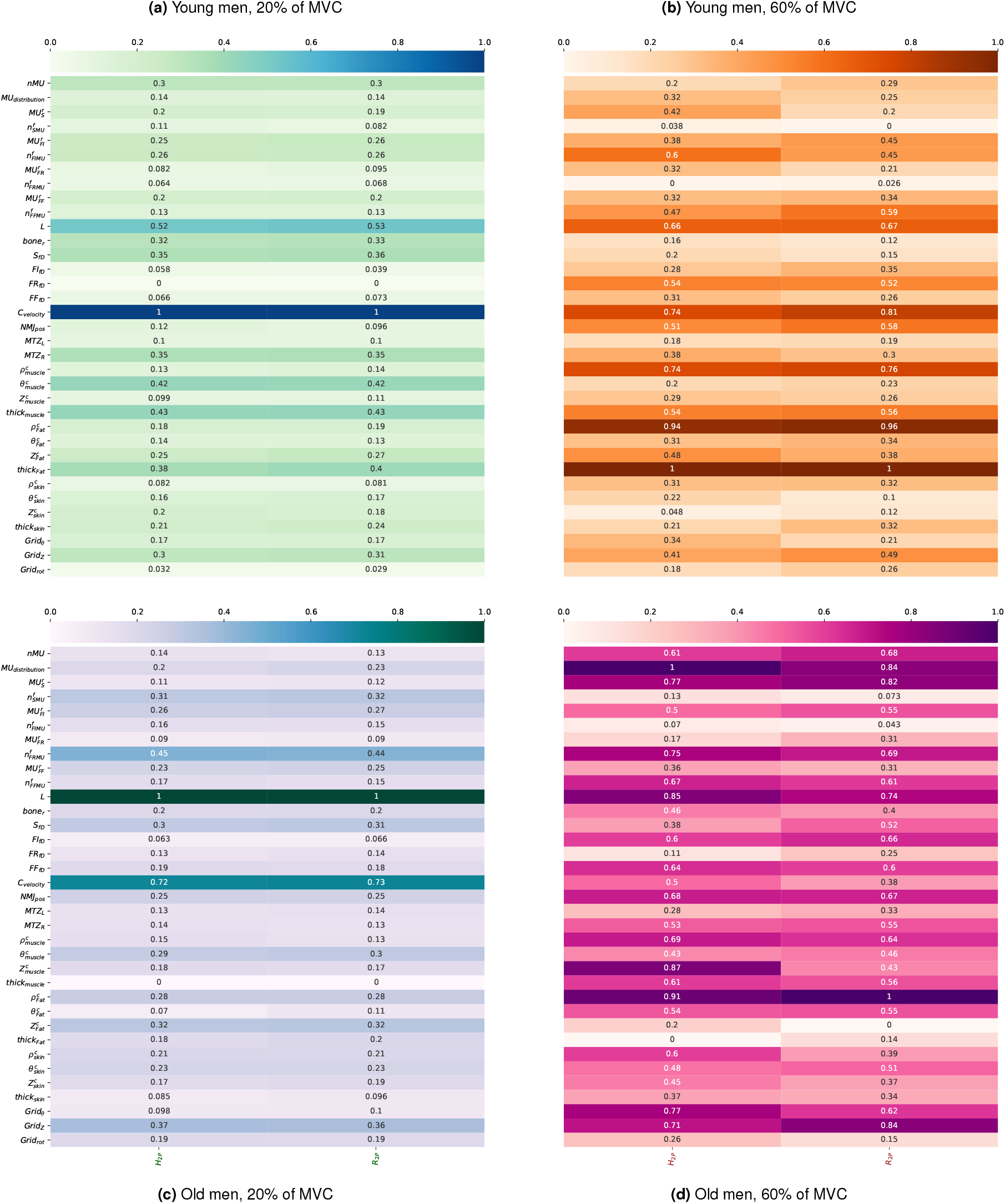
Impacts of model inputs on extracted amlitude HD-sEMG Bi-variate features (RMSM results). RMSM impact indice = Normalized *SI* (from 0 (light color) to 1 (dark color)). Ranking of neuromuscular inputs according to age (young (a,b) and old men (c,d)), and level of force contractions (low (a,c) and high(b,d)).

##### group 1: MDF and MNF

These two features are useful as fatigue indicators. Its relation to conduction velocity of fibers was proved as linear impact on HD-sEMG signals at isometric contractions. The length of fibers (*L*) is in the second position, and the rest of inputs have a negligible effects. These two inputs have a monotonic effect with tendency of *C*_*velocity*_ to have an almost linear effect (RMSM screening).

##### group 2: SM1, SM2 and SM3

The *C*_*velocity* has the most important impact. At LC, the muscle thickness and conductivity have the same important impact for YM and OM. This influence decrease for higher order of SM. Then a reduced group of parameters have an intermediate influence. This group changes slightly one or more of its members when changing contraction level or age. We observe that this group contains muscle thickness (*thick*_*muscle*_) and conductivities 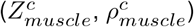. For (YM, HC), this group contains, in addition, the number of slow fibers per MU. For (OM, LC), this group contains in addition the number of slow fibers per MU. Four parameters share the first ranks: conduction velocity, muscle thickness, longitudinal and radial muscle conductivities respectively. We note that, with RMSM screening, these features have an identical distribution of parameters, and that the influential parameters are located in the “almost-monotonic” zone.

##### group 3: TTP and MNP

The most influential parameters for all categories are: the conduction velocity of fibers (*C*_*velocity*_), the longitudinal conductivity of fibers 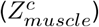, and the muscle thickness (*thick*_*muscle*_). For LC categories, the number of slow fibers per MU has a large impact (FF fibers at HC). We observe that the MU repartition have an important impact for OM at HC.

##### group 3: Freq_ratio and VDF

All parameters have a negligible effect, expecting *conduction*_*velocity*.

##### group 3: Peak_Freq

Most of parameters have a significant impact.

### F. Bi-variate features

Two features are selected: the linear correlation coefficient: Pearson coefficient, and the non-linear coefficient: *H*_2_*P*. We cannot observe a significant difference between YM and OM. The changes are observed between LC and HC. For LC, only two parameters enhance correlation between recording system channels: *L* and *C*_*velocity*_. For HC, much more parameters have a large impact (almost of parameters).

### G. Features/Parameters selections

The features selected by elimination on the VIF criterion were the same for both young and elder men, only the contraction level induced a slightly different results. At the low contraction level, **AAC** and **ARV** are the only remaining features under the threshold value of 10. At the higher level, **DASDV** is then also selected. This small subset of features are the ones with the least colinearity and can then be used first to select our global subset of import parameters but also later on to build cost function for parameter identification.

Then, using this subset of features, the sensitivity analysis results are fed to scoring methods, a majority of them agree on the following parameters as globally the most important within the model:

#### low contraction

*boneR*, CV, L, MUSnbFibers, nUM, sigRho@0, sigRho@1, sigTheta@0, sigZ@0, thick@0

#### high contraction

*boneR*, centery, CV, MUFFnbFibers, MUrepartition, MUSfibersDiameter, nUM, sigRho@1, sigZ@0, thick@0

In italics is the radius of the bone, which shows an importance only in the case of older men, but then is always ranked first.

As expected, at low contraction level the slow fibers are important whereas the fast ones dominate at higher contraction levels. Conduction velocity, number of motor units and parameters of the conductor volumes are presents in all cases. At high contraction, the lateral position of the recording system rise in importance, most likely due to the higher spatial heterogeneity of the HD-sEMG signals due to the firing of fast motor units closer to the skin.

## Discussions

The purpose of this study is the identification of anatomical and neural muscle factors with the largest impact on the HD-sEMG signals simulated with our model. The application of RMSM method on the HD-sEMG signals generated by the neuromuscular analytic model allows the identification of parameters with small and large impact for young and old men at low and high contractions. The muscle anatomy for each age groups is taken from literature, e.g. number of fibers, cross sectional area,etc.

In this study, we have demonstrated that amplitude features of simulated HD-sEMG signals are sensitive to: muscle thickness, the conduction velocity, and muscle conductivities at LC. In addition, at HC, HD-sEMG is impacted by the MU number and the electrode grid position. Many studies Del Vecchio et al. (2017); Dideriksen et al. (2011); Farina et al. (2004) showed that EMG amplitude is correlated to the increase of force exerted due to the increase of MU recruited. The conduction velocity impact at high contractions level is mentioned in Del Vecchio et al. (2018) and clearly observed in Fig. 7 for YM and OM at HC. The importance of electrode placement was evoked in many previous studies Campanini et al. (2007); Mesin et al. (2009), due to its high dependency to source locations, innervation zone, and the non-homogeneity of the conductor volume. This study shows that the orientation and position of recording system have a large influence at HC for HD-sEMG signals with amplitude features. However, in Campanini et al. (2007); Mesin et al. (2009), it was not mentioned that this effect increases with high contractions. In fact, in the modeling approach depicted in Carriou et al. (2016), fast sources (FF MU) are positioned at muscle surface layer and much more recruited at HC. Which can explain the large effect of recording system position at HC. This sensitivity analysis shows a high influence of slow and fast fibers/MU number at low and high contractions respectively on HD-sEMG signals. This result is consistent with finding in Lee et al. (2013); Rainoldi (2008); Wakeling et al. (2006). These studies demonstrated that for LC only slow fibers/MU are recruited to exert force. For HC, fast fibers are recruited after recruiting slower ones. Their number is determinant in defining muscle capacity for maintaining force.

This study states that fiber diameters have not a direct impact on the HD-sEMG signal compared to number of fibers. The loss of fibers and MU diameters with aging does not have a large impact on the simulated HDsEMG signals. This statement should be viewed cautiously. In fact, it can indicate that apoptosis phenomena have much more impact on the force generated by muscle than other phenomena such as the atrophy for example. From such conclusion, we can imply that the decline of muscle functional with aging can be related mainly to the neural system (denervation-reinnervation failure) Hepple and Rice (2016).

However, other relevant observation with RMSM results should be mentioned: the muscle thickness have more impact with aged subjects for amplitude features (Fig. 7). This parameter is related to muscle cross section area CSA, and its size reduction is due to both apoptosis and atrophy. From such result, the previous statement is not totally effective. Moreover, the conduction velocity of fibers, which have a large impact in this study, is a function of fiber diameters as depicted in Carriou et al. (2016). Therefore, the HD-sEMG can be considered as very sensitive to atrophy but indirectly, through muscle thickness and conduction velocity. This is more consistent with many studies correlating the reduction of muscle size to the loss of force with aging.

All These expected results add to the credibility the modeling approaches used in the neuromuscular model we used.

For frequency domain features, it was obvious that the conduction velocity of fibers and the muscle conductivity have the most important impacts. This is in agreement with experimental results Marco et al. (2017); Barandun et al. (2009) and can validate assumptions and approaches applied in Carriou et al. (2016). With RMSM results, we have observed no significant difference between YM and OM at LC and HC. This can be related to two facts: (i) FD features are useful to detect muscle fatigue and this aspect is not considered in Carriou et al. (2016), and (ii) this model is not incorporating phenomena related to muscle aging.

However, the importance of RMSM results can be extended to cover other important sides. One of the purposes of applying a sensitivity analysis method on HD-sEMG signals is to develop a patient personalized model to evaluate and diagnosis muscle aging. A parameter values identification with inverse methods is contemplated to reach a reliable diagnosis. RMSM results offer a map of relationships between inputs and outputs. This will facilitate the decision of which muscle factor should be identified and by which feature and at which condition (age, contractions level).

Other important aspect can be underlined from RMSM results: a homogenization of the most influential muscle parameters for subjects, if possible. In fact, this homogenization can be valuable and crucial for reliable clinical assessment of muscle force and diseases. For a reliable assessment of Neuropathies, for example, it is better to neutralize the effect of muscle thickness on the EMG signals by selecting subjects with a small variation scale for this parameter. Commonly, clinical test practitioners are not enough cautioned against this factor. Usually, studies for clinical EMG measures use essentially the body mass index (BMI) as the main criteria to select subjects/patients. Moreover, this study shows that the position electrodes can be important for many features and at some contractions level (Fig. 7). A rotation of the recording grid by few degrees, or a translation by few millimeters can impact greatly the measured HD-sEMG signals. Therefore, this study suggested more care for this kind of aspects when evaluating muscle diseases and health by EMG techniques.

It was verified in many clinical studies that, for time domain features, parameters related to the structure and morphology of muscle have a large impact on the measured EMG signals. The identification of important and negligible parameters is a supporting argument for the validation of modeling approaches and assumptions applied and depicted in Carriou et al. (2016, 2018).

## Conclusions

In this study, a complete global sensitivy analisys of a multiscale HD-sEMG muscle was performed correlating to anatomical and neural properties of the neuromuscular system, within the specific context of muscle aging. For this purpose, muscle aging modifications have been incorporated in the model parameters and tested in term of influence on classical HD-sEMG descriptors. From the results, it appears that key parameters modified by aging have a important influence and must took into account carefully for next studies combining experimental and simulated data. These results illustrated neuromuscular modeling as a valuable tool to evaluate health and disease muscle state with low cost and in reduced time. However, for a reliable evaluation of muscle aging, modeling approaches should be enhanced to better describe structural, morphological, and functional age-related phenomena.

## ACKNOWLEDGEMENTS

This work was carried out and funded in the framework of the Labex MS2T. We thank equally the Region Hauts-de-France for co-funding this work.

## Bibliography

Barandun, M., von Tscharner, V., Meuli-Simmen, C., Bowen, V., and Valderrabano, V. (2009). Frequency and conduction velocity analysis of the abductor pollicis brevis muscle during early fatigue. Journal of Electromyography and Kinesiology, 19(1):65–74. doi: 10.1016/j.jelekin.2007.07.003.

Boccia, G., Dardanello, D., Rosso, V., Pizzigalli, L., and Rainoldi, A. (2015). The Application of sEMG in Aging: A Mini Review. Gerontology, 61(5):477–484. doi: 10.1159/000368655.

Campanini, I., Merlo, A., Degola, P., Merletti, R., Vezzosi, G., and Farina, D. (2007). Effect of electrode location on EMG signal envelope in leg muscles during gait. Journal of Electromyography and Kinesiology, 17(4):515–526. doi: 10.1016/j.jelekin.2006.06.001.

Campolongo, F., Cariboni, J., and Saltelli, A. (2007). An effective screening design for sensitivity analysis of large models. Environmental Modelling & Software, 22(10):1509–1518. doi: 10.1016/j.envsoft.2006.10.004.

Carriou, V., Boudaoud, S., Laforet, J., and Ayachi, F. S. (2016). Fast generation model of high density surface EMG signals in a cylindrical conductor volume. Computers in Biology and Medicine, 74:54–68. doi: 10.1016/j.compbiomed.2016.04.019.

Carriou, V., Boudaoud, S., and Laforet, J. (2018). Speedup computation of HD-sEMG signals using a motor unit-specific electrical source model. Medical & Biological Engineering & Computing, 56(8):1459–1473. doi: 10.1007/s11517-018-1784-5.

Del Vecchio, A., Negro, F., Felici, F., and Farina, D. (2017). Associations between motor unit action potential parameters and surface EMG features. Journal of Applied Physiology (Bethesda, Md.: 1985), 123(4):835–843. doi: 10.1152/japplphysiol.00482.2017.

Del Vecchio, A., Bazzucchi, I., and Felici, F. (2018). Variability of estimates of muscle fiber conduction velocity and surface EMG amplitude across subjects and processing intervals. Journal of Electromyography and Kinesiology, 40:102–109. doi: 10.1016/j.jelekin.2018.04.010.

Dideriksen, J. L., Enoka, R. M., and Farina, D. (2011). Neuromuscular adjustments that constrain submaximal EMG amplitude at task failure of sustained isometric contractions. Journal of Applied Physiology (Bethesda, Md.: 1985), 111(2):485–494. doi: 10.1152/japplphysiol.00186.2011.

Douania, I., Laforêt, J., and Boudaoud, S. (2023). Robust morris screening method (rmsm) for complex physiological models. Computer Methods and Programs in Biomedicine, 231:107368. doi: 10.1016/j.cmpb.2023.107368.

Farina, D. (2008). Counterpoint: Spectral properties of the surface emg do not provide information about motor unit recruitment and muscle fiber type. Journal of Applied Physiology, 105(5): 1673–1674. doi: 10.1152/japplphysiol.90598.2008a.

Farina, D., Merletti, R., and Enoka, R. M. (2004). The extraction of neural strategies from the surface EMG. Journal of Applied Physiology (Bethesda, Md.: 1985), 96(4):1486–1495. doi: 10.1152/japplphysiol.01070.2003.

González-Izal, M., Malanda, A., Gorostiaga, E., and Izquierdo, M. (2012). Electromyographic models to assess muscle fatigue. Journal of Electromyography and Kinesiology, 22(4):501–512. doi: 10.1016/j.jelekin.2012.02.019.

Hepple, R. T. and Rice, C. L. (2016). Innervation and neuromuscular control in ageing skeletal muscle. The Journal of Physiology, 594(8):1965–1978. doi: 10.1113/JP270561.

Larsson, L. (1978). Morphological and functional characteristics of the ageing skeletal muscle in man. A cross-sectional study. Acta Physiologica Scandinavica. Supplementum, 457:1–36.

Lee, S. S. M., de Boef Miara, M., Arnold, A. S., Biewener, A. A., and Wakeling, J. M. (2013). Recruitment of faster motor units is associated with greater rates of fascicle strain and rapid changes in muscle force during locomotion. The Journal of Experimental Biology, 216(2):198–207. doi: 10.1242/jeb.072637.

Lexell, J., Henriksson-Larsén, K., Winblad, B., and Sjöström, M. (1983). Distribution of different fiber types in human skeletal muscles: Effects of aging studied in whole muscle cross sections. Muscle & Nerve, 6(8):588–595. doi: 10.1002/mus.880060809.

Macaluso, A. and De Vito, G. (2004). Muscle strength, power and adaptations to resistance training in older people. European Journal of Applied Physiology, 91(4):450–472. doi: 10.1007/s00421-003-0991-3.

Marco, G., Alberto, B., and Taian, V. (2017). Surface EMG and muscle fatigue: multi-channel approaches to the study of myoelectric manifestations of muscle fatigue. Physiological Measurement, 38(5):R27–R60. doi: 10.1088/1361-6579/aa60b9.

Mesin, L., Merletti, R., and Rainoldi, A. (2009). Surface EMG: The issue of electrode location. Journal of Electromyography and Kinesiology, 19(5):719–726. doi: 10.1016/j.jelekin.2008.07.006.

Morris, M. D. (1991). Factorial Sampling Plans for Preliminary Computational Experiments. Technometrics, 33(2):161–174. doi: 10.2307/1269043.

Nair, K. S. (2005). Aging muscle. The American Journal of Clinical Nutrition, 81(5):953–963. doi: 10.1093/ajcn/81.5.953.

Rainoldi, A. (2008). Spectral properties of the surface EMG can characterize/do not provide information about motor unit recruitment strategies and muscle fiber type. Journal of Applied Physiology (Bethesda, Md.: 1985), 105(5):1678.

Sjöström, M., Lexell, J., and Downham, D. Y. (1992). Differences in fiber number and fiber type proportion within fascicles. A quantitative morphological study of whole vastus lateralis muscle from childhood to old age. The Anatomical Record, 234(2):183–189. doi: 10.1002/ar.1092340205.

Sobol, I. M. (2001). Global sensitivity indices for nonlinear mathematical models and their Monte Carlo estimates. Mathematics and Computers in Simulation, 55(1):271–280. doi: 10.1016/S0378-4754(00)00270-6.

Tieland, M., Trouwborst, I., and Clark, B. C. (2018). Skeletal muscle performance and ageing. Journal of Cachexia, Sarcopenia and Muscle, 9(1):3–19. doi: 10.1002/jcsm.12238.

Verdijk, L. B., Koopman, R., Schaart, G., Meijer, K., Savelberg, H. H. C. M., and van Loon, L. J. C. (2007). Satellite cell content is specifically reduced in type II skeletal muscle fibers in the elderly. American Journal of Physiology. Endocrinology and Metabolism, 292(1):E151–157. doi: 10.1152/ajpendo.00278.2006.

Wakeling, J. M., Uehli, K., and Rozitis, A. I. (2006). Muscle fibre recruitment can respond to the mechanics of the muscle contraction. Journal of the Royal Society Interface, 3(9):533–544. doi: 10.1098/rsif.2006.0113.

